# The pharmacokinetics and pharmacodynamics of 4-methylumbelliferone and its glucuronide metabolite in mice

**DOI:** 10.1101/2022.08.18.504417

**Authors:** Nadine Nagy, Gernot Kaber, Naomi L. Haddock, Aviv Hargil, Jayakumar Rajadas, Sanjay V. Malhotra, Marc A. Unger, Adam R. Frymoyer, Paul L. Bollyky

**Author notes:** Corresponding Author: Nadine Nagy, PhD, Division of Infectious Diseases and Geographic Medicine, Department of Medicine, Stanford University School of Medicine, Beckman B237, 279 Campus Drive, Stanford, CA 94305-5107, USA.

## Abstract

Hyaluronan (HA) is an extracellular matrix glycosaminoglycan, with important roles in chronic inflammation, cancer and autoimmunity. 4-methylumbelliferone (4-MU), a small molecule inhibitor of HA synthases, is widely used to study HAs interactions with the surrounding tissues and the immune cells. There is substantial experimental and therapeutic interest in using oral 4-MU to inhibit HA synthesis, but pharmacokinetic and pharmacodynamic data on treatment routes have been lacking. Moreover, it recently became clear that the main metabolite of 4-MU, 4-methlyumbelliferyl glucuronide (4-MUG), is bioactive. We therefore sought to define the pharmacokinetics and pharmacodynamics of 4-MU and its active metabolite 4-MUG in mice. Single dose mouse studies showed that 4-MU administered intravenously (i.v.) resulted in 100-fold higher 4-MU exposure compared to oral (p.o.) administration. The 4-MU ratio AUC i.v./AUC p.o. was 96/1. 4-MUG exposures were much higher than 4-MU exposures after both 4-MU i.v. and p.o. administration, but only small differences in 4-MUG exposure were seen after 4-MU i.v. versus p.o. administration. The 4-MUG metabolite was also administered as a single dose both i.v. and p.o. and showed a 25.9% bioavailability. Compared to 4-MUG p.o. dosing, 1.14 higher 4-MUG exposures were seen after 4-MU p.o. dosing. 4-MU exposure after 4-MUG p.o. administration was minimal but similar to 4-MU exposure after 4-MU p.o. administration. In mice treated for several weeks with 4-MU in chow, the 4-MU concentration immediately drops after treatment was stopped, whereas the 4-MUG concentration showed a peak 1 hour after treatment stop. In a build-up study, 4-MU and 4-MUG treatment in mice lead to a plateau of 4-MU concentration starting at 4 days post treatment start. These 4-MU and 4-MUG concentration findings *in vivo* will inform future clinical studies and experimental work with 4-MU.

## Introduction

4-methylumbelliferone (4-MU) is a small molecule inhibitor of hyaluronan (HA) synthesis (1), which has been shown to inhibit HA production in multiple cell lines and tissue types *in vitro* and *in vivo* (2,3,4). HA is an extracellular matrix glycosaminoglycan with many roles in tissue function and development (5-7), which has been identified as a driving factor in inflammation (2,8,9). Greatly elevated HA levels have been found in chronically inflamed tissues (10-12), the tumor microenvironment, in fibrosis, and at sites of autoimmunity (13,14). There is substantial experimental and therapeutic interest in inhibiting HA synthesis. The mechanism of action by which 4-MU inhibits HA synthesis has been described (15). UDP-GlcUA and UDP-GlcNAc are the substrates of the HA synthesis, their availability thereby limits HA synthesis. 4-MU mainly functions by depleting the HA precursor UDP-GlcUA by activation of UDP-glucuronyl transferases (UGTs) (15) and 4-MU also downregulates the HAS expression (16).

4-MU is a commercially available drug in Europe and Asia and is approved for use in humans to prevent biliary spasm (17-20). Several clinical trials in humans have been published on 4-MU, all demonstrated excellent safety data (21-27). Right now, 4-MU is under investigation in human clinical trials as a safety study at multiple doses, and the treatment for HA-associated fibrotic liver and autoimmune biliary diseases (ClinicalTrials.gov Identifiers: NCT00225537, NCT02780752). The biggest downside to 4-MU are its poor pharmacokinetics, which are thought to limit its use outside the biliary tract. The use of 4-MU in other diseases at other sites of the body is desirable due to HA’s involvement in many different chronic and acute diseases. The systemic oral bioavailability of 4-MU is reported to be < 3%, mostly due to extensive first pass glucuronidation in the liver and small intestine (18,28). If first pass metabolism is bypassed by giving the dose intravenously (i.v.), the systemic exposure achieved can be more than 10-30 fold higher than after the same dose given orally (p.o.) (29). However, due to the high clearance of 4-MU, systemic concentrations will decrease rapidly after an i.v. dose and peripheral exposures will likely be almost undetectable by 4 to 6 hours after a 4-MU dose. The pharmacodynamic (PD) and pharmacokinetic (PK) properties of p.o. vs i.v. 4-MU administration are of great importance and will be investigated in this study.

Any 4-MU that reaches the systemic circulation is rapidly metabolized with a half-life of 28 minutes in humans and <1% of a given dose is excreted unchanged in the urine (18,28). 4-MUs half-life in mice is with 3 minutes (min.) even shorter. Consequently, the median plasma concentration of the 4-MU metabolite, 4-methlyumbelliferyl glucuronide (4-MUG), is more than 3,000-fold higher than that of 4-MU in mouse models (1). Analogous findings have been reported in healthy human volunteers (28). Metabolism of the drug occurs via conjugation to either a glucuronic (4-MUG), or a sulfate (4-MUS). The glucuronide (4-MUG) is the predominant pathway and accounts for over 90% of its metabolism (29,30). 4-MUG is more hydrophilic and gets eliminated in the bile and urine (30). Biliary eliminated 4-MUG likely undergoes further enterohepatic recirculation with reabsorption of the metabolite from the intestine and ultimate elimination in the urine via the kidney. This is supported by a healthy volunteer pharmacokinetic study in which 93% of a single i.v. dose of 4-MU was eliminated as the 4-MUG metabolite in the urine (29). The PK and PD properties of 4-MU and 4-MUG will be investigated in this study.

Despite the poor bioavailability and a short half-life, oral administration of 4-MU nonetheless inhibits HA synthesis *in vivo*, and has demonstrated a reduction of severity and frequency in different chronic and autoimmune diseases (1,9,14). Future animal studies and clinical studies would benefit from a more thorough understanding of the pharmacokinetics and pharmacodynamics of 4-MU and the 4-MUG metabolite to help support dosing and target exposure. Furthermore, to our knowledge PK/PD data for multiple dosing with 4-MU and 4-MUG have not been reported previously. Here, we have used mass spectrometry to interrogate the pharmacokinetics and pharmacodynamics of 4-MU and 4-MUG in mice with single and multiple dosing.

## Materials and Methods

### Mice

All animals were bred and maintained under specific pathogen-free conditions, with free access to food and water, in the animal facilities at Bioduro (Beijing, China) and Stanford University Medical School (Stanford, CA). Male CD-1 mice were bred in house at Bioduro, and male C57Bl/6J mice were purchased from The Jackson Laboratories (Bar Harbor, Me).

### 4-MU and 4-MUG treatment

4-MU (Alfa Aesar) was pressed into the mouse chow by TestDiet^®^ and irradiated before shipment, as previously described (4). We previously determined that this chow formulation delivers 250 mg/mouse/day, yielding a serum drug concentration of 640.3 ± 17.2 nmol/L in mice, as measured by HPLC-MS. 4-MUG (ChemImpex) was distributed in the drinking water at a concentration of 2 mg/ml, delivering 10 mg/mouse/day, yielding a serum drug concentration of 357.1 ± 72.6 ng/mL in mice, as measured by LC-MS/MS. Mice were initiated on 4-MU and 4-MUG at eight weeks of age, unless otherwise noted, and were maintained on this diet until they were euthanized, unless otherwise noted.

### Pharmacokinetic Study in CD-1 mice for 4-MU

Male CD-1 mice, 7-9 weeks of age, were given 4-MU and 4-MUG at 25 mg/kg i.v. at a vehicle concentration of 5 mg/mL in an injection volume of 5 mL/kg. 10% DMSO (Sigma-Aldrich and Tedia)/ 50%PEG400 (Fluka)/ 40% water was used as vehicle. Mice were fed at the time of injection. Blood samples were collected at 2 min, 5 min, 15 min, 30 min, 1 hour (hr), 2 hr, 4 hr, 6 hr, 8 hr and 24 hr from the saphenous vein, temporally put on ice until plasma was processed. Plasma samples were stored at −80C until analysis.

CD-1 mice were given 4-MU at 25 mg/kg p.o. via gavage at a vehicle concentration of 2.5 mg/mL in a volume of 10 mL/kg. 5% methyl cellulose (Sigma) was used as vehicle. Mice were fed at the time of experiment. Blood samples were collected at 5 min, 15 min, 30 min, 1 hr, 2 hr, 4 hr, and 6 hr, from the saphenous vein, temporally put on ice until plasma was processed. Plasma samples were stored at −80C until analysis.

The bioanalysis was performed via LC/MS/MS (AB Science API 5500 and AB Science Triple Quad 6500+) using the Analyst Software 1.7.1 a Shimadzu pump (LC-20AD) and Acquity Ultra Performance LC System (Waters) and a Kinetex column (2.6u C18 100A 30 mm x 3 mm) and an Aquity UPLC BEH C18 2.1*50 mm column (Waters). 50 ng/mL Tolbutamide in methanol/acetonitrile (1:1, v/v) was used as internal standard.

### Caco-2 permeability assessment

Caco-2 cells were cultured to confluency, trypsinised and seeded onto filter transwell inserts at a density of 32K cells/well in DMEM cell culture medium (Gibco). Following an overnight attachment period, the cell medium was replaced every other day. The cell monolayer was ready for assessment 22 days post seeding. A 25 mM HBSS (Sigma)/HEPES (Gibco) solution pH 7.4 and a HBSS/HEPES 0.1% BSA (VWR Life Science) solution were prepared. 10 uM 4-MU (Sigma) and 4-MUG (Sigma) working solutions were prepared as well as internal standards and quenching solutions (25/50 ng/mL Terfenadine (Sigma)/Tolbutamide. The TEER (Millicell ERS-2) value of each well was measured, the wells were only used when their TEER values were greater than 600 Ohms/cm^2^. The cell plates were prepared as follows, the cell culture medium from the basolateral side (B) and apical side (A) were aspirated. For A to B direction, the donor plate A is was rinsed with 400 ul HBSS/Hepes, the recipient plate B was rinsed with 400 ul HBSS/Hepes/0.1% BSA. For the B to A direction, the donor plate B was rinsed with 1200 ul HBSS/Hepes solution and the recipient plate A was rinsed with 400 ul HBSS/Hepes/0.1% BSA. The plates were dosed as follows, for the A to B direction, 500 ul of the compound or control working solution was added to the donor plate A. 1300 ul of HBSS/Hepes/0.1 BSA solution was added to the recipient plate B. This was vice versa for the B to A direction. Plates were pre-incubated for 10 min and 100 ul sample was collected as o min sample for future analysis. The cell plates were incubated for another 90 min and samples were taken after that. The samples were prepared for LC-MS/MS analysis. The MS detection was performed using Sciex API 4500 Q Trap and 5500 instruments. Each compound was analyzed by reverse phase HPLC.

### Take away study

4-MU (Alfa Aesar) was pressed into the mouse chow by TestDiet^®^ and irradiated before shipment, as previously described (4). We previously determined that this chow formulation delivers 250 mg/mouse/day. 4-MUG (ChemImpex) was distributed in the drinking water at a concentration of 2 mg/ml, delivering 10 mg/mouse/day. Male C57Bl/6J mice were treated with 5% 4-MU (Alpha Aesar) and 2 mg/ml 4-MUG (Sigma) starting at 8 weeks of age and maintained on this diet until they were euthanized, unless otherwise noted. The mice were treated for 5 weeks. At the end of the 5 weeks of treatment the mice were transferred to clean cages with regular chow and blood samples were taken at T0, T1, T2, T6, T12 and T24 hours and processed to plasma. Plasma samples were stored at −80C for further analysis.

### Built up study

Male C57Bl/6J mice were treated with 5% 4-MU (Alpha Aesar) and 2 mg/ml 4-MUG (Sigma) starting at 8 weeks of age for 20 days. During the time of treatment blood samples were taken from the mice at T0, T1, T4, T7, T15 and T20 days after treatment start. Plasma samples were stored at −80C for further analysis.

### Liquid Chromatography Tandem Mass Spectrometry (LC-MS/MS) analysis of 4-MU and 4-MUG concentrations in mouse serum

4-MU and 4-MUG concentrations were measured via mass spectrometry. 4-methylumbelliferone-13C4 (Toronto Research Chemicals, Ontario, Canada) was used as the IS for 4-MU and 7-hydroxycoumarin β-D-glucuronide (Toronto Research Chemicals, Ontario, Canada) as the IS for 4-MUG. The neat stock solutions of 4-MU and 4-MUG were mixed and diluted in 50% methanol to prepare the spiking solutions ranging from 1 ng/mL to 5000 ng/mL for each compound.

For calibration standards, 25 µl of blank serum was mixed with 25 µl of the spiking solutions. For samples to be tested, 25 µl of serum was mixed with 25 µl of 50% methanol to make up the volume. 25 µl of a mixture of the two IS (1000 ng/ml each in 50% methanol) was then added. After vortexing all standards and samples, 150 µl of methanol/acetonitrile 20:80 (v/v) was added to the mixture and the sample was further vortexed vigorously for 1 min followed by centrifugation at 3,000 rpm for 10 min. 100 µl of the supernatant was taken and diluted with 200µl of Milli Q water.

The LC-MS/MS system consists of an AB SCIEX QTRAP 4000 mass spectrometer linked to a Shimadzu UFLC system. Mobile phase A is HPLC grade water. Mobile phase B is HPLC grade acetonitrile. LC separation was carried out on a Phenomenex Luna PFP(2) column (3 µm, 150 × 2 mm) with isocratic elution using 45% mobile phase B and a flow rate of 0.4 ml/min at room temperature. The analysis time was 2.5 min. 10 µl of the extracted sample was injected. The mass spectrometer was operated in the negative mode with the following multiple-reaction monitoring (MRM) transitions: m/z 174.7→132.9 for 4-MU, m/z 178.7→134.9 for 4-MU-13C4 (IS), m/z 350.8→174.9 for 4-MUG and m/z 336.9→160.9 for 7-hydroxy coumarin β-D-glucuronide (IS). Data acquisition and analysis were performed using the Analyst 1.6.1 software (AB SCIEX).

### Pharmacokinetic Analysis

The C_max_ (maximum plasma concentration) and T_max_ (time of maximum concentration) were obtained directly from the observed data. The terminal rate constant (λ_z_) was determined by linear regression analysis of the terminal portion of the log plasma concentration–time curve. The terminal half-life (t_1/2_) was calculated as ln 2/λ_z_. AUC_0-last_ was found using the linear/logarithmic trapezoidal method. Summation of AUC_last_ and the concentration at the last measured point divided by *λ*_z_ yielded AUC_0-∞_. Bioavailability (F) was calculated as AUC_0-∞,iv_ / AUC_0-∞,po_. To allow direct comparisons of AUC_0-∞_ across 4MU and 4MUG dose formulations, AUC_0-∞_ after 4MUG dose was dose-adjusted for molar equivalent of 4MU using a factor of 2. The mean residence time (MRT) was calculated as the ratio of the area under the first moment curve (AUMC_0–∞_) divided by AUC_0–∞_ minus the estimated mean absorption time (MAT) (i.e., MRT=(AUMC_0–∞_ / AUC_0–∞_) - MAT). MAT was the reciprocal of the first-order absorption rate (k_a_) constant estimated when fitting concentration data to a two-compartmental model with first-order absorption.

### Statistical analysis

Data are expressed as means ± SEM of *n* independent measurements. A *p* value of <0.05 was considered significant. Significance of the difference between the means of two or three groups of data was evaluated using a two-tailed *t* test or one-way ANOVA with Šidák’s multiple comparisons post-test. GraphPad Prism 8.1.2 software was used to perform data analysis.

## Results

### 4-MU administration i.v. and p.o., 4-MU measured

To establish the basic PK properties of 4-MU in mice, a single dose of 25 mg/kg 4-MU was administered i.v. and p.o. (**Table 1, 2**). The mean terminal half-life (t_1/2_) of 4-MU in plasma was 0.182 hr for i.v. and 0.346 hr for p.o. administration. The initial plasma drug concentration at time zero following the bolus i.v. injection (C_0_) was 21852 ng/mL. The time to reach maximal plasma concentration (t_max_) following p.o. administration was 0.139 hr and the maximal plasma 4-MU concentration (C_max_) was 100 ng/mL. The area under the plasma concentration-time curve extrapolated from time t to infinity as a percentage of total AUC (AUC_last_) was 2875 hr*ng/mL for i.v. and 25.9 hr*ng/mL for p.o. administration. Similarly, the area under the plasma concentration-time curve from time zero to infinity (AUC_Inf_) was 2876 hr*ng/mL for i.v and 27.8 hr*ng/mL for p.o. administration. The mean infinite residence time (MRT_Inf_) for 4-MU after i.v. administration was 0.0866 hr and 0.451 hr for p.o. administration. The dose-normalized area under the plasma concentration-time curve from time 0 to infinity (AUC_Inf_/D) for 4-MU administered p.o. was 1.11 hr*kg*ng/mL/mg. The systemic bioavailability (F) for the p.o. administered 4-MU dose was 0.967 %. For the 4-MU administration i.v. the apparent volume of distribution during the terminal phase (V_Z_) was 2.31 L/kg, and the apparent volume of distribution at steady state (V_SS_) was 0.757 L/kg. The apparent total body clearance of the drug from plasma (CL) was 145 mL/min/kg for i.v. 4-MU administration.

**Table 1:** Basic PK properties of 4-MU in mice.

| Animal | Treatment | Administration | measured | t<sub>1/2</sub> (hr) | C<sub>0</sub> (ng/mL) | AUC<sub>last</sub> (hr*ng/mL) | AUC<sub>inf</sub> (hr*ng/mL) | AUC<sub>extr</sub> (%) | V<sub>z</sub> (L/kg) | V<sub>ss</sub> (L/kg) | CL (mL/min/kg) | MRT<sub>inf</sub> (hr) |
| --- | --- | --- | --- | --- | --- | --- | --- | --- | --- | --- | --- | --- |
| Mouse 1 | 4-MU (25mg/kg) | i.v. | 4-MU | 0.315 | 20195 | 2818 | 2820 | 0.0607 | 4.04 | 0.942 | 148 | 0.106 |
| Mouse 2 | 4-MU (25mg/kg) | i.v. | 4-MU | 0.118 | 26334 | 3111 | 3113 | 0.0584 | 1.36 | 0.607 | 134 | 0.0756 |
| Mouse 3 | 4-MU (25mg/kg) | i.v. | 4-MU | 0.114 | 19027 | 2695 | 2696 | 0.0485 | 1.53 | 0.722 | 155 | 0.0779 |
| Mean | 4-MU (25mg/kg) | i.v. | 4-MU | 0.182 | 21852 | 2875 | 2876 | 0.0559 | 2.31 | 0.757 | 145 | 0.0866 |
| SD | 4-MU (25mg/kg) | i.v. | 4-MU | 0.115 | 3925 | 214 | 214 | 0.0065 | 1.50 | 0.170 | 11 | 0.0171 |

**Table 2:** Basic PK properties of 4-MU in mice.

| Animal | Treatment | Administration | measured | t<sub>1/2</sub> (hr) | t<sub>max</sub> (hr) | C<sub>max</sub> (ng/mL) | AUC<sub>last</sub> (hr*ng/mL) | AUC<sub>inf</sub> (hr*ng/mL) | AUC<sub>extr</sub> (%) | MRT<sub>inf</sub> (hr) | AUC<sub>inf</sub>/D (hr*kg*ng/mL/mg) | F (%) |
| --- | --- | --- | --- | --- | --- | --- | --- | --- | --- | --- | --- | --- |
| Mouse 4 | 4-MU (25mg/kg) | p.o. | 4-MU | 0.405 | 0.25 | 31.3 | 19.5 | 20.1 | 2.99 | 0.52 | 0.804 | 0.699 |
| Mouse 5 | 4-MU (25mg/kg) | p.o. | 4-MU | 0.446 | 0.083 | 55.3 | 17.7 | 22.4 | 21.1 | 0.62 | 0.896 | 0.779 |
| Mouse 6 | 4-MU (25mg/kg) | p.o. | 4-MU | 0.188 | 0.083 | 214 | 40.4 | 40.9 | 1.28 | 0.21 | 1.64 | 1.42 |
| Mean | 4-MU (25mg/kg) | p.o. | 4-MU | 0.346 | 0.139 | 100 | 25.9 | 27.8 | 8.47 | 0.451 | 1.11 | 0.967 |
| SD | 4-MU (25mg/kg) | p.o. | 4-MU | 0.139 | 0.096 | 99.3 | 12.6 | 11.4 | 11.0 | 0.216 | 0.46 | 0.397 |

### 4-MU administration i.v. and p.o., 4-MUG measured

In the same experiment as described above, 4-MU’s major metabolite 4-MUG was assessed after 25 mg/kg 4-MU was administered i.v. and p.o. (**Table 3, 4**). The mean half-life (t_1/2_) of 4-MUG in the serum was 1.29 hr for i.v. and 3.79 hr for p.o. administration. The initial plasma 4-MUG concentration at time zero following the bolus i.v. injection (C_0_) was 25400 ng/mL. The time to reach maximal plasma concentration following p.o. administration (t_max_) was 0.25 hr and the maximal plasma 4-MU concentration (C_max_) was 28733 ng/mL. The area under the plasma concentration-time curve extrapolated from time t to infinity as a percentage of total AUC (AUC_last_) was 45389 hr*ng/mL for i.v. and 20009 hr*ng/mL for p.o. administration. The area under the plasma concentration-time curve from time zero to infinity (AUC_Inf_) was 45842 hr*ng/mL for i.v and 27086 hr*ng/mL for p.o. administration. The area under the plasma concentration-time curve extrapolated from time to infinity as a percentage of total AUC (AUC_Extr_) was 1.01% for i.v. and 17.6% for p.o. 4-MU administration. The mean infinite residence time (MRT_Inf_) for 4-MU after i.v. administration was 0.638 hr and 4.000 hr for p.o. administration.

**Table 3:** Assessment of 4-MUG properties after 4-MU administration.

| Animal | Administration | Administration | measured | t<sub>1/2</sub> (hr) | C<sub>0</sub> (ng/mL) | AUC<sub>last</sub> (hr*ng/mL) | AUC<sub>inf</sub> (hr*ng/mL) | AUC<sub>extr</sub> (%) | MRT<sub>inf</sub> (hr) |
| --- | --- | --- | --- | --- | --- | --- | --- | --- | --- |
| Mouse 1 | 4-MU (25mg/kg) | i.v. | 4-MUG | 1.46 | 16100 | 54570 | 54982 | 0.751 | 0.611 |
| Mouse 2 | 4-MU (25mg/kg) | i.v. | 4-MUG | 0.961 | 30200 | 41093 | 41194 | 0.246 | 0.503 |
| Mouse 3 | 4-MU (25mg/kg) | i.v. | 4-MUG | 1.44 | 29900 | 40505 | 41351 | 2.05 | 0.800 |
| Mean | 4-MU (25mg/kg) | i.v. | 4-MUG | 1.29 | 25400 | 45389 | 45842 | 1.01 | 0.638 |
| SD | 4-MU (25mg/kg) | i.v. | 4-MUG | 0.28 | 8055 | 7956 | 7916 | 0.93 | 0.150 |

**Table 4:** Assessment of 4-MUG properties after 4-MU administration.

| Animal | Administration | Administration | measured | t<sub>1/2</sub> (hr) | t<sub>max</sub> (hr) | C<sub>max</sub> (ng/mL) | AUC<sub>last</sub> (hr*ng/mL) | AUC<sub>inf</sub> (hr*ng/mL) | AUC<sub>extr</sub> (%) | MRT<sub>inf</sub> (hr) |
| --- | --- | --- | --- | --- | --- | --- | --- | --- | --- | --- |
| Mouse 1 | 4-MU (25mg/kg) | p.o. | 4-MUG | 1.72 | 0.25 | 22100 | 18470 | 18911 | 2.33 | 1.10 |
| Mouse 2 | 4-MU (25mg/kg) | p.o. | 4-MUG | 8.94 | 0.25 | 20900 | 20490 | 41259 | 50.3 | 10.4 |
| Mouse 3 | 4-MU (25mg/kg) | p.o. | 4-MUG | 0.705 | 0.25 | 43200 | 21067 | 21089 | 0.105 | 0.55 |
| Mean | 4-MU (25mg/kg) | p.o. | 4-MUG | 3.79 | 0.25 | 28733 | 20009 | 27086 | 17.6 | 4.00 |
| SD | 4-MU (25mg/kg) | p.o. | 4-MUG | 4.49 | 0.00 | 12543 | 1363 | 12323 | 28.4 | 5.51 |

### 4-MU and 4-MUG concentration in mice over time after 4-MU i.v. and p.o. administration

4-MU was administered to mice at 25 mg/kg i.v. and p.o., and subsequent the serum concentration of 4-MU and 4-MUG was measured over time (**Figure 1**). The mice dosed 4-MU i.v started at 0.033 hr with 17800 ng/mL 4-MU whereas the mice dosed 4-MU p.o. starting at 0.083 hr had a mean 4-MU concentration of 100 ng/mL (**Figure 1A-C**). At 0.5 hr the 4-MU concentration after i.v. administration went down to 72.7 ng/mL and 16.7 ng/mL was measured for p.o. administration (**Figure 1A-C**). At the 2 hr timepoint 4-MU was barely detectable in the serum independent of route of administration, 1.25 ng/mL was measured for i.v. and 0.34 ng/mL for p.o. (**Figure 1A-C**).

**Figure 1.**
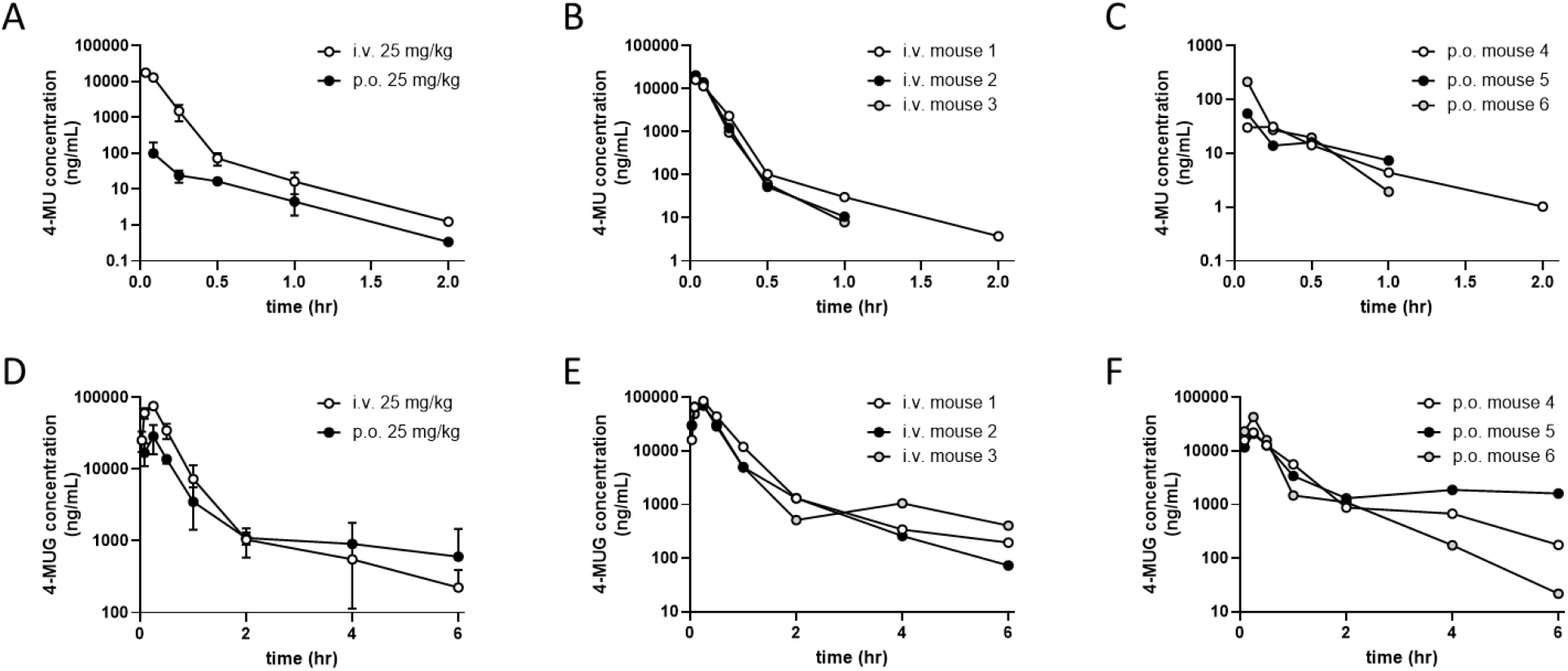
4-MU i.v. and p.o administration. CD-1 mice, 7-9 weeks of age, were given 4-MU i.v. and p.o. Blood samples were collected at 2 min, 5 min, 15 min, 30 min, 1 hr, 2 hr, 4 hr, 6 hr, 8 hr and 24 hr. The 4-MU and 4-MUG concentration in the blood samples was determined via LC-MS/MS. **A**. Mean 4-MU concentration after i.v and p.o. administration at different timepoints. **B**. 4-MU concentration after i.v. administration, individual animals. **C**. 4-MU concentration after p.o. administration, individual animals. **D**. Mean 4-MUG concentration after i.v. and p.o. administration at different timepoints. **E**. 4-MUG concentration after i.v. administration, individual animals. **F**. 4-MUG concentration after p.o. administration, individual animals. N = 3 mice per group.

After 4-MU administration, 4-MUG was detected at 0.033 hr at 25400 ng/mL in the i.v. group, compared to 16933 ng/mL at 0.083 hr for the p.o. administration group (**Figure 1D-F**). 1 hour after 4-MU administration, the 4-MUG concentration in the mouse serum after i.v. and p.o. administration was almost identical with 1045 ng/mL for i.v. and 1110 ng/mL for p.o. (**Figure 1D-F**). At the last measurement time point of 6 hr the 4-MUG concentration after 4-MU i.v. injection was 225 ng/mL and 603 ng/mL for p.o. administration (**Figure 1D-F**).

### 4-MUG administration i.v. and p.o., 4-MUG measured

To establish the basic PK properties of 4-MUG in mice, 25 mg/kg 4-MUG, the major metabolite of 4-MU, was administered i.v. and p.o. (**Table 5, 6**). The mean half-life (t_1/2_) of 4-MUG in the serum was 3.17 hr for i.v. and 1.14 hr for p.o. administration. The initial plasma drug concentration at time zero (C_0_) following the bolus i.v. injection was 102477 ng/mL. The time to reach maximal plasma concentration (t_max_) following p.o. administration was 3.33 hr and the maximal plasma 4-MUG concentration (C_max_) was 2800 ng/mL. The area under the plasma concentration-time curve extrapolated from time t to infinity as a percentage of total AUC (AUC_last_) was 33783 hr*ng/mL for i.v. and 8506 hr*ng/mL for p.o. administration. The area under the plasma concentration-time curve from time zero to infinity (AUC_Inf_) was 33792 hr*ng/mL for i.v and 8766 hr*ng/mL for p.o. administration. The area under the plasma concentration-time curve extrapolated from time to infinity as a percentage of total AUC (AUC_Extr_) was 0.0262% for i.v. and 2.91% for p.o. 4-MU administration. The mean infinite residence time (MRT_Inf_) for 4-MUG after i.v. administration was 0.524 hr and 3.46 hr for p.o. administration. The dose-normalized area under the plasma concentration-time curve from time 0 to infinity (AUC_Inf_/D) for 4-MUG administered p.o. was 351 hr*kg*ng/mL/mg. The systemic availability (F) for the p.o. administered 4-MUG dose was 25.9%. For the 4-MUG administration i.v. the apparent volume of distribution during the terminal phase (V_Z_) was 0.681 L/kg, and the apparent volume of distribution at steady state (V_SS_) was 0.0805 L/kg. The apparent total body clearance of the drug from plasma (CL) was 2.51 mL/min/kg for i.v. 4-MUG administration.

**Table 5:** Basic properties of 4-MUG in mice.

| Animal | Treatment | Administration | measured | t<sub>1/2</sub> (hr) | C<sub>0</sub> (ng/mL) | AUC<sub>last</sub> (hr*ng/mL) | AUC<sub>inf</sub> (hr*ng/mL) | AUC<sub>extr</sub> (%) | V<sub>z</sub> (L/kg) | V<sub>ss</sub> (L/kg) | CL (mL/min/kg) | MRT<sub>inf</sub> (hr) |
| --- | --- | --- | --- | --- | --- | --- | --- | --- | --- | --- | --- | --- |
| Mouse 1 | 4-MUG (25mg/kg) | i.v. | 4-MUG | 3.09 | 90225 | 31708 | 31717 | 0.0267 | 0.704 | 0.0820 | 2.63 | 0.520 |
| Mouse 2 | 4-MUG (25mg/kg) | i.v. | 4-MUG | 3.46 | 123807 | 39900 | 39907 | 0.0192 | 0.625 | 0.0501 | 2.09 | 0.400 |
| Mouse 3 | 4-MUG (25mg/kg) | i.v. | 4-MUG | 2.95 | 93398 | 29741 | 29751 | 0.0325 | 0.716 | 0.109 | 2.80 | 0.650 |
| Mean | 4-MUG (25mg/kg) | i.v. | 4-MUG | 3.17 | 102477 | 33783 | 33792 | 0.0262 | 0.681 | 0.0805 | 2.51 | 0.524 |
| SD | 4-MUG (25mg/kg) | i.v. | 4-MUG | 0.26 | 18540 | 5388 | 5387 | 0.0066 | 0.050 | 0.0296 | 0.37 | 0.125 |

**Table 6:** Basic properties of 4-MUG in mice.

| Animal | Treatment | Administration | measured | t<sub>1/2</sub> (hr) | t<sub>max</sub> (hr) | C<sub>max</sub> (ng/mL) | AUC<sub>last</sub> (hr*ng/mL) | AUC<sub>inf</sub> (hr*ng/mL) | AUC<sub>extr</sub> (%) | MRT<sub>inf</sub> (hr) | AUC<sub>inf</sub>/D (hr*kg*ng/mL/mg) | F (%) |
| --- | --- | --- | --- | --- | --- | --- | --- | --- | --- | --- | --- | --- |
| Mouse 4 | 4-MUG (25mg/kg) | p.o. | 4-MUG | 1.04 | 4 | 2960 | 10023 | 10330 | 2.97 | 4.14 | 413 | 30.6 |
| Mouse 5 | 4-MUG (25mg/kg) | p.o. | 4-MUG | 1.43 | 2 | 3970 | 8135 | 8494 | 4.24 | 2.94 | 340 | 25.1 |
| Mouse 6 | 4-MUG (25mg/kg) | p.o. | 4-MUG | 0.959 | 4 | 1470 | 7362 | 7475 | 1.51 | 3.30 | 299 | 22.1 |
| Mean | 4-MUG (25mg/kg) | p.o. | 4-MUG | 1.14 | 3.33 | 2800 | 8506 | 8766 | 2.91 | 3.46 | 351 | 25.9 |
| SD | 4-MUG (25mg/kg) | p.o. | 4-MUG | 0.25 | 1.15 | 1258 | 1369 | 1447 | 1.36 | 0.61 | 58 | 4.3 |

### 4-MUG administration i.v. and p.o., 4-MU measured

In the same experiment to establish the basic PK properties of 4-MUG, 4-MU was assessed after 25 mg/kg 4-MUG was administered i.v. and p.o. (**Table 7,8**). The mean half-life (t_1/2_) of 4-MU in the serum was 0.265 hr for i.v. and 0.00 for p.o. administration. The initial plasma 4-MUG concentration at time zero (C_0_) following the bolus i.v. injection was 504 ng/mL. The time to reach maximal plasma concentration (t_max_) following p.o. administration was 0.27 hr and the maximal plasma 4-MU concentration (C_max_) was 10.2 ng/mL. The area under the plasma concentration-time curve extrapolated from time t to infinity as a percentage of total AUC (AUC_last_) was 218 hr*ng/mL for i.v. and 14.3 hr*ng/mL for p.o. administration. The area under the plasma concentration-time curve from time zero to infinity (AUC_Inf_) was 219 hr*ng/mL for i.v and 0.00 hr*ng/mL for p.o. administration. The area under the plasma concentration-time curve extrapolated from time to infinity as a percentage of total AUC (AUC_Extr_) was 0.579% for i.v. and 0.00% for 4-MU measured after p.o. 4-MUG administration. The mean infinite residence time (MRT_Inf_) for 4-MU after i.v. administration of 4-MUG was 0.082 hr and 0.00 hr for p.o. administration.

**Table 7:**
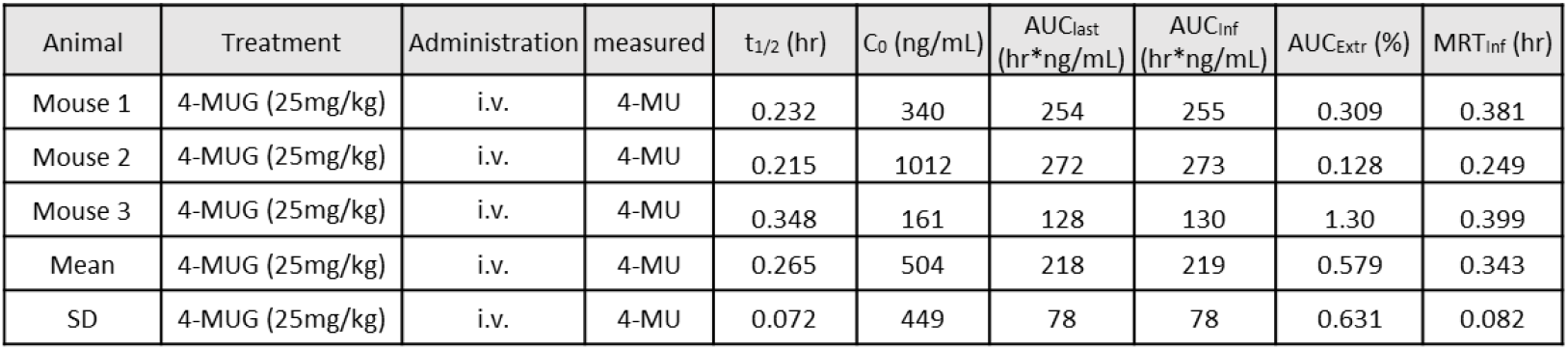
Assessment of 4-MU properties after 4-MUG administration.

| Animal | Treatment | Administration | measured | t<sub>1/2</sub> (hr) | C<sub>0</sub> (ng/mL) | AUC<sub>last</sub> (hr*ng/mL) | AUC<sub>inf</sub> (hr*ng/mL) | AUC<sub>extr</sub> (%) | MRT<sub>inf</sub> (hr) |
| --- | --- | --- | --- | --- | --- | --- | --- | --- | --- |
| Mouse 1 | 4-MUG (25mg/kg) | i.v. | 4-MU | 0.232 | 340 | 254 | 255 | 0.309 | 0.381 |
| Mouse 2 | 4-MUG (25mg/kg) | i.v. | 4-MU | 0.215 | 1012 | 272 | 273 | 0.128 | 0.249 |
| Mouse 3 | 4-MUG (25mg/kg) | i.v. | 4-MU | 0.348 | 161 | 128 | 130 | 1.30 | 0.399 |
| Mean | 4-MUG (25mg/kg) | i.v. | 4-MU | 0.265 | 504 | 218 | 219 | 0.579 | 0.343 |
| SD | 4-MUG (25mg/kg) | i.v. | 4-MU | 0.072 | 449 | 78 | 78 | 0.631 | 0.082 |

**Table 8:** Assessment of 4-MU properties after 4-MUG administration.

| Animal | Treatment | Administration | measured | t<sub>1/2</sub> (hr) | t<sub>max</sub> (hr) | C<sub>max</sub> (ng/mL) | AUC<sub>last</sub> (hr*ng/mL) | AUC<sub>inf</sub> (hr*ng/mL) | AUC<sub>extr</sub> (%) | MRT<sub>inf</sub> (hr) |
| --- | --- | --- | --- | --- | --- | --- | --- | --- | --- | --- |
| Mouse 1 | 4-MUG (25mg/kg) | p.o. | 4-MU | 0.00 | 4 | 4.96 | 7.73 | 0.00 | 0.00 | 0.00 |
| Mouse 2 | 4-MUG (25mg/kg) | p.o. | 4-MU | 0.00 | 2 | 19.2 | 19.2 | 0.00 | 0.00 | 0.00 |
| Mouse 3 | 4-MUG (25mg/kg) | p.o. | 4-MU | 0.00 | 2 | 6.37 | 16.0 | 0.00 | 0.00 | 0.00 |
| Mean | 4-MUG (25mg/kg) | p.o. | 4-MU | 0.00 | 2.67 | 10.2 | 14.3 | 0.00 | 0.00 | 0.00 |
| SD | 4-MUG (25mg/kg) | p.o. | 4-MU | NA | 1.15 | 7.8 | 5.9 | NA | NA | NA |

### 4-MUG and 4-MU concentration in mice over time after 4-MUG i.v. and p.o. administration

4-MUG was administered to mice at 25 mg/kg i.v. and p.o., the serum concentration of 4-MUG and 4-MU was measured (**Figure 2**). The mice dosed i.v started at 0.083 hr with 74133 ng/mL 4-MUG whereas the mice dosed p.o. starting at 0.25 hr had a mean 4-MUG concentration of 209 ng/mL (**Figure 2A-C**). At 1 hr the 4-MUG concentration after i.v. administration went down to 3590 ng/mL and the 4-MUG concentration after 1 hr p.o. administration went up to 359 ng/mL (**Figure 2A-C**). At the 8 hr timepoint 4-MUG was barely detectable in the serum after i.v. administration, with 28.1 ng/mL compared to 154 ng/mL still detectable after 8hr after p.o. administration of 4-MUG (**Figure 2A-C**).

**Figure 2.**
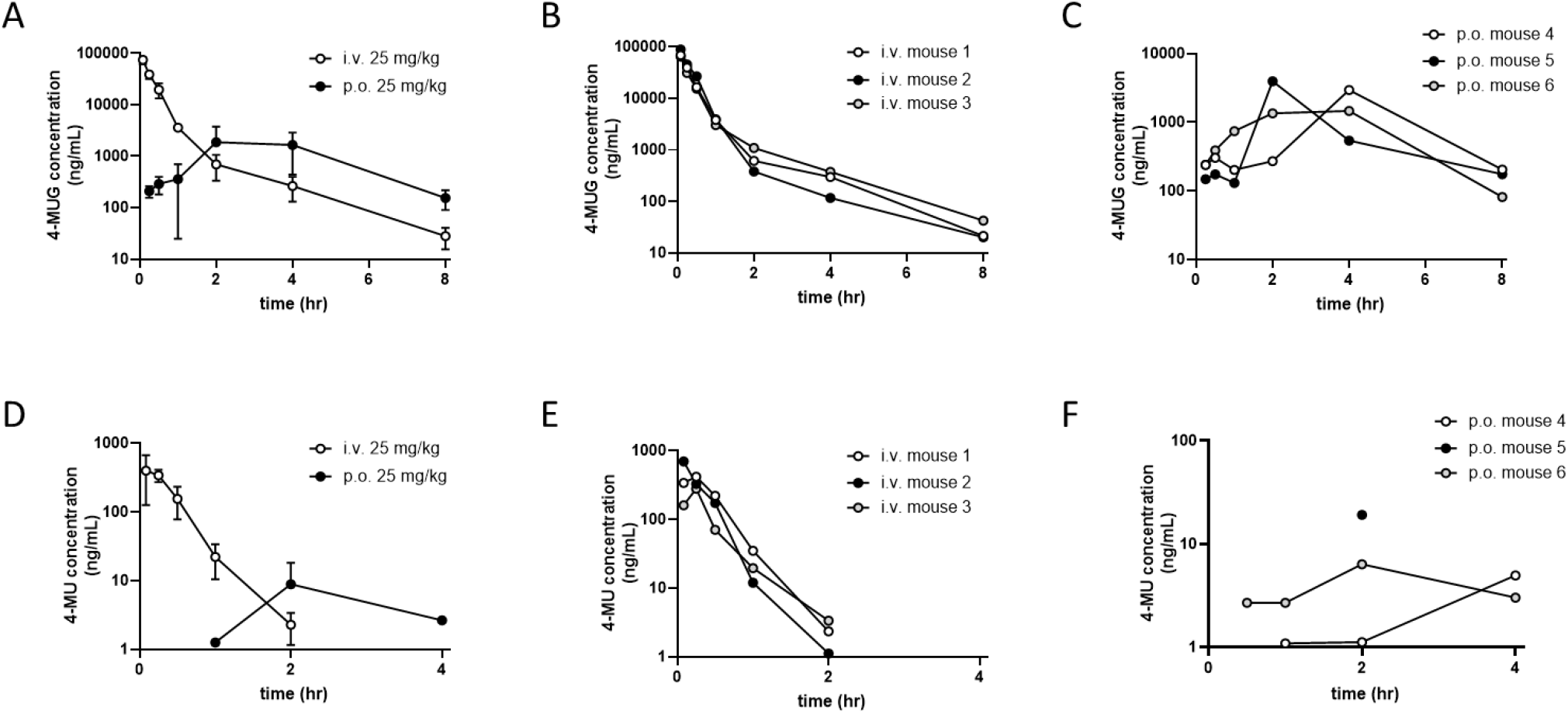
4-MUG i.v. and p.o administration. CD-1 mice, 7-9 weeks of age, were given 4-MUG i.v. and p.o. Blood samples were collected at 2 min, 5 min, 15 min, 30 min, 1 hr, 2 hr, 4 hr, 6 hr, 8 hr and 24 hr. The 4-MU and 4-MUG concentration in the blood samples was determined via LC-MS/MS. **A**. Mean 4-MUG concentration after i.v and p.o. administration at different timepoints. **B**. 4-MUG concentration after i.v. administration, individual animals. **C**. 4-MUG concentration after p.o. administration, individual animals. **D**. Mean 4-MU concentration after i.v. and p.o. administration at different timepoints. **E**. 4-MU concentration after i.v. administration, individual animals. **F**. 4-MU concentration after p.o. administration, individual animals. N = 3 mice per group.

After 4-MUG administration, 4-MU was detected at 0.083 hr at 399 ng/mL in the i.v. group, compared to 0.00 ng/mL for the p.o. administration group (**Figure 2D-F**). 1 hour after 4-MUG administration, the 4-MU concentration in the mouse serum after i.v. was 22.2 ng/mL and p.o. barely detectable with 1.26 ng/mL (**Figure 2D-F**). At the last measurement time point with usable data at 4 hr the 4-MU concentration after 4-MUG i.v. injection was not detectable and 2.66 ng/mL for p.o. administration (**Figure 2D-F**).

### 4-MU and 4-MUG permeability assessment

In an Caco-2 permeability experiment we assessed the permeability and Pgp substrate classification for 4-MU and 4-MUG (**Table 9**). For 4-MU the permeability was classified as high and the 4-MUG permeability was classified as low, both substances were not classified as Pgp substrates. The % recovery from A → B and from B → A was between 86.9 and 93.4 for 4-MU and 104 and 103 for 4-MUG. The parameter apparent permeability Papp from A → B and from B → A was 42.9 and 37.2×10^−6^ cm/s for 4-MU and 0.156 and 0.129 ×10^−6^ cm/s for 4-MUG. The efflux ratio was 0.865 for 4-MU and 0.823 for 4-MUG.

**Table 9:** Caco-2 permeability assessment of 4-MU and 4-MUG.

| Compound ID | % Recovery (N = 2) |  | Papp (x10<sup>-6</sup> cm/s) (N = 2) |  | Efflux ratio | Permeability Classification | Pgp substrate |
| --- | --- | --- | --- | --- | --- | --- | --- |
|  | A → B | B → A | A → B | B → A |  |  |  |
| 4-MU | 86.9 | 93.3 | 42.9 | 37.2 | 0.865 | high | NO |
| 4-MUG | 104 | 103 | 0.156 | 0.129 | 0.823 | low | NO |
| atenolol | 103 | 93.4 | 0.300 | 0.402 | 1.34 | low | NO |
| propanolol | 90.1 | 91.4 | 29.1 | 19.6 | 0.671 | high | NO |
| taxel | 108 | 96.0 | 0.944 | 27.4 | 29.0 | low | YES |

### 4-MU and 4-MUG treatment stop study

To evaluate how fast after treatment stop the 4-MU and 4-MUG plasma concentration decreases, we treated mice for 5 weeks with 4-MU and 4-MUG and then stopped the treatment (**Figure 3**). We took blood samples at different timepoints after treatment stop up to 24 hrs. We found that the 4-MU concentration after 4-MU treatment stop significantly decreased after 1 hour from 4960 ng/mL to 1156 ng/mL (**Figure 3A**). At 2, 6 and 12 hours after treatment stop the 4-MU concentration was 460 ng/mL, 220 ng/mL and 92 ng/mL (**Figure 3A**). At 24 hours after treatment stop the 4-MU concentration in the serum was close to background noise with 4.4 ng/mL (**Figure 3A**). In the same 4-MU treated animals we measured the 4-MUG concentration in the serum after treatment stop (**Figure 3B**). Here we saw a slightly different drug concentration profile. Immediately after treatment stop the 4-MUG concentration in the serum was 220333 ng/mL and increased to 354667 ng/mL 1 hour after treatment stop (**Figure 3B**). This increase was followed by a rapid decrease to 31133 ng/mL at 2 hours after treatment stop and declined further at 6 and 12 hours were 4-MUG concentrations of 1461 ng/mL and 520 ng/mL were measured (**Figure 3B**). 24 hours after 4-MU treatment stop the 4-MUG concentration was with 140 ng/mL at the level of background noise (**Figure 3B**).

**Figure 3.**
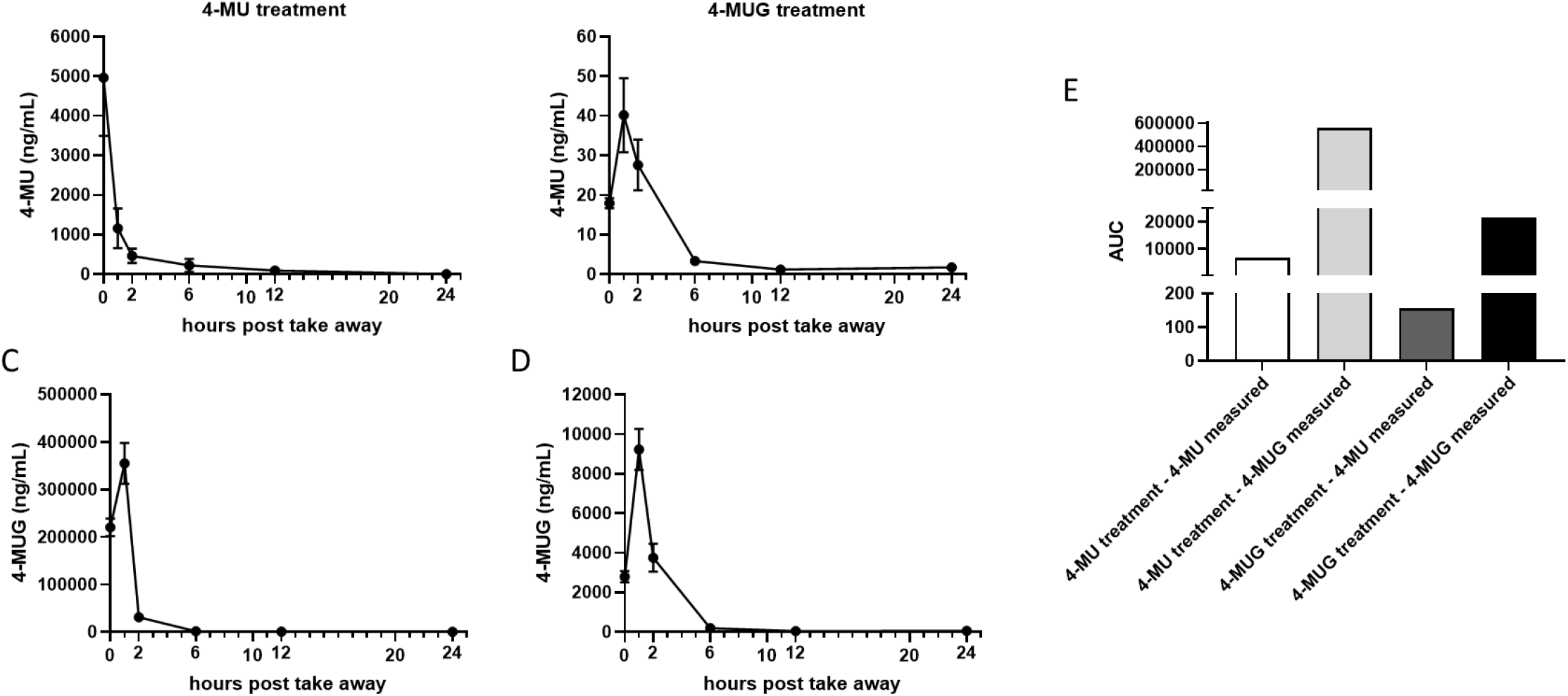
4-MU and 4-MUG treatment stop study. C57Bl/6J mice were treated with 4-MU and 4-MUG starting at 8 weeks of age. The mice were treated for 5 weeks, and at the end of 5 weeks the treatment was stopped and blood samples were taken at T0, T1, T2, T6, T12 and T24 hours. The 4-MU and 4-MUG concentration in the blood samples was determined via LC-MS/MS. **A**. 4-MU treated mice, 4-MU measured. **B**. 4-MUG treated mice, 4-MU measured. **C**. 4-MU treated mice, 4-MUG measured. **D**. 4-MUG treated mice, 4-MUG measured. **E**. AUC for all treatment groups. N = 5 mice per group.

In the 4-MUG treatment group we found after treatment stop very low levels of 4-MU (**Figure 3C**). At treatment stop the 4-MU concentration was 17 ng/mL, increased to 40 ng/mL 1 hour after treatment stop and declined to background noise at 6, 12 and 24 hours ranging between 3 and 1 ng/mL (**Figure 3C**). We saw much higher concentrations of 4-MUG after 4-MG treatment stop (**Figure 3D**). At the time of 4-MUG treatment stop, the 4-MUG concentration was 2796 ng/mL, which increased to 9223 ng/mL 1 hour after treatment stop (**Figure 3D**). 2 hours after treatment stop the 4-MUG concentration declined to 3752ng/mL and at 6 hours after treatment stop the 4-MUG concentration declined further almost to background level to 199 ng/mL, followed by 44 ng/mL and 57 ng/mL at the 12 hour and 24 hour time points (**Figure 3D**). The AUC calculations of this experiment revealed a bigger AUC when 4-MUG was measured, after initial 4-MUG treatment, the highest AUC was calculated for 4-MUG measurement when the initial treatment was 4-MU (**Figure 3E**).

### MU and 4-MUG treatment built up study

To evaluate how fast and to what concentrations 4-MU and 4-MUG would built up in the plasma, we treated mice for 20 days with 4-MU and 4-MUG (**Figure 4**). We took blood samples at different timepoints from treatment start up to 20 days later. We found that the 4-MU concentration after 4-MU treatment reached a plateau after 4 days of treatment (**Figure 4A**). One day after treatment start a 4-MU concentration of 541 ng/mL was detected, which increased to 1371 ng/mL at 4 days after treatment start (Fig. 4A). The 4-MU measurement at 7 days and 15 days post treatment start showed with 1228 ng/mL and 1516 ng/mL similar concentration level (**Figure 4A**). We noticed a decline in 4-MU concentration to 931 ng/mL at day 20 post treatment start (**Figure 4A**). In the same 4-MU treated animals we measured the 4-MUG concentration from treatment start and saw the same plateau starting at day 4, with a slight increase at day 15 (**Figure 4B**). The 4-MUG concentration after 1 day of 4-MU treatment was 27228 ng/mL, which increased to 262400 ng/mL 4 days after treatment start (**Figure 4B**). The 4-MUG concentrations at day 7 and day 20 after treatment start were with 297800 ng/mL and 268540 ng/mL very similar (**Figure 4B**). An increase in 4-MUG concentration could be detected at 15 days post treatment start, here the concentration reached 386460 ng/mL (**Figure 4B**).

**Figure 4.**
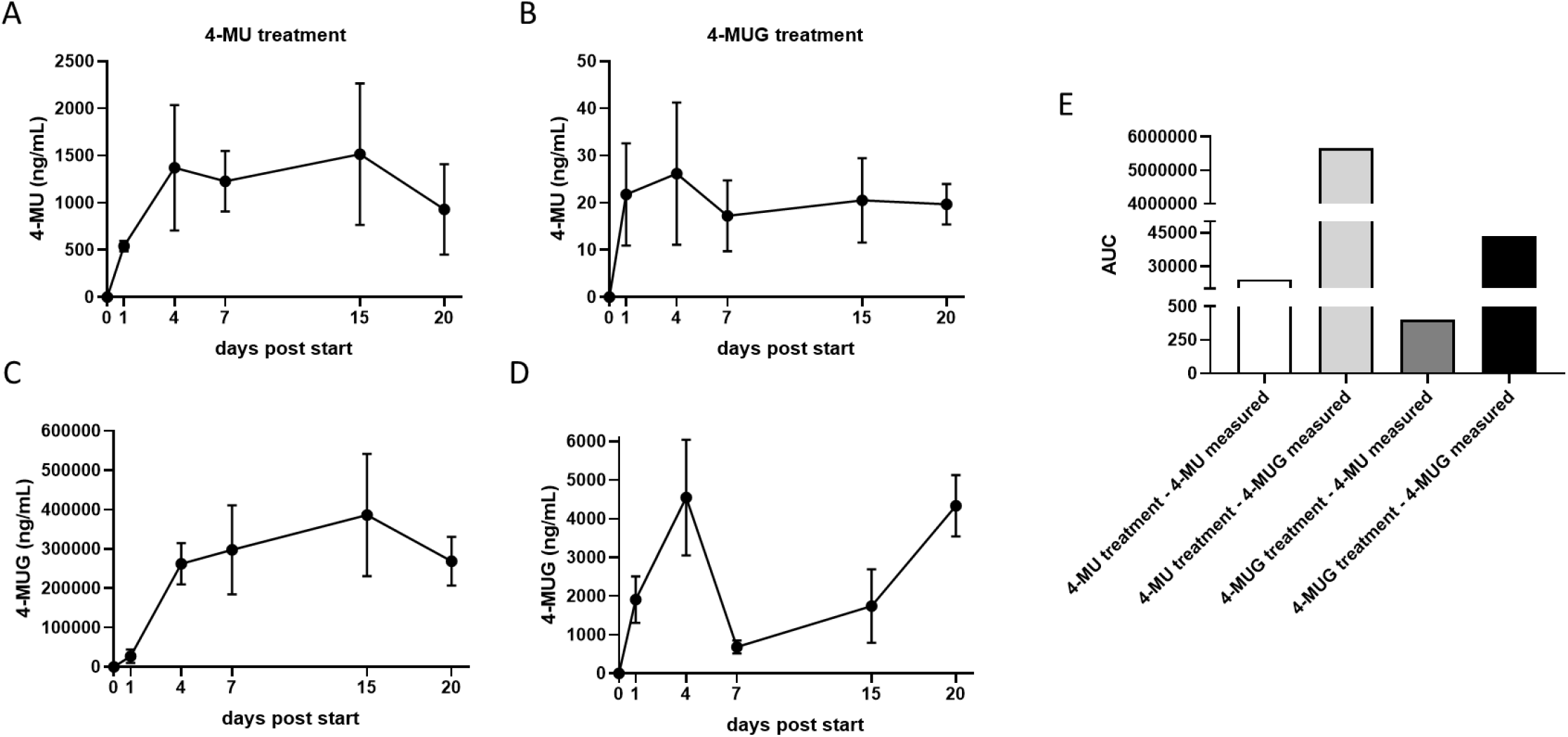
4-MU and 4-MUG treatment built up study. C57Bl/6J mice were treated with 4-MU and 4-MUG starting at 8 weeks of age for 20 days. During the time of treatment blood samples were taken from the mice at T0, T1, T4, T7, T15 and T20 days after treatment start. The 4-MU and 4-MUG concentration in the blood samples was determined via LC-MS/MS. **A**. 4-MU treated mice, 4-MU measured. **B**. 4-MUG treated mice, 4-MU measured. **C**. 4-MU treated mice, 4-MUG measured. **D**. 4-MUG treated mice, 4-MUG measured. **E**. AUC for all treatment groups. N = 5 mice per group.

In the 4-MUG treatment group the 4-MU concentration was in general very low, here a plateau was reached after 1 day of treatment (**Figure 4C**). One day post treatment start 4-MU reached concentrations of 21 ng/mL, which increased to 26 ng/mL at day 4 post treatment (**Figure 4C**). The 4-MU concentrations after 4-MUG treatment at days 7, 15 and 20 post treatment start where with 17 ng/mL, 20 ng/mL and 19 ng/mL quite similar (**Figure 4C**). When we looked at the 4-MUG concentration after 4-MUG treatment, we found a massive drop at day 7 post treatment start followed by another increase at day 20 (**Figure 4D**). 1 day after treatment start the 4-MUG concentration was at 1907 ng/mL and increased further to 4546 ng/mL 4 days after treatment start (**Figure 4D**). This was followed by a huge decrease in 4-MUG concentration at day 7 post treatment start to 685 ng/mL (**Figure 4D**). The 4-MUG concentration after 4-MUG treatment increased again at day 7 and day 20 post treatment start to 1740 ng/mL and 4332 ng/mL (**Figure 4D**). The AUC calculations of this experiment revealed a bigger AUC when 4-MUG was measured, after initial 4-MUG treatment, the highest AUC was calculated for 4-MUG measurement when the initial treatment was 4-MU (**Figure 4E**).

## Discussion

In these studies, we have examined the PK for both oral and i.v. 4-MU and 4-MUG in mice when given as single as well as multiple dosing protocols.

We find that the most of the circulating drug present *in vivo* exists as 4-MUG. This is consistent with previous PK studies in animals that demonstrated the extraction of 4-MU by the gastrointestinal system (pre-hepatic) to be ∼40% and extraction by the liver as high as 97% (20) due to the major hepatic and intestinal UGTs involved in drug metabolism (31). As a result of this high extraction, the fraction of an administered oral dose of hymecromone that reaches the systemic circulation (post-hepatic) as unchanged drug (i.e. the bioavailability) is very low. These data indicate that It is important to understand the 4-MUG PK during oral 4-MU therapy, given the expected much higher exposures of the metabolite relative to the parent in peripheral tissues.

In our studies comparing p.o. and i.v. delivery of 4-MU, we report that the half-life of 4-MU given as a single dose via i.v. administration (0.182 hr) was doubled when given p.o. (0.346 hr). The half-life of the resulting 4-MUG metabolite given as a single dose i.v. (1.29 hr) was almost tripled when given p.o. (3.79 hr) was one third. Comparing p.o. and i.v. delivery of 4-MUG, we report the half-life was almost three times higher for i.v. (3.17 hr) administration compared to p.o. (1.14 hr). When we dosed 4-MUG and measured 4-MU, only a low half-life for i.v. administration could be observed, the oral administration did not yield in measurable 4-MU concentrations. Consistent with this, in a Caco-2 experiment we found that the permeability of 4-MU is high and the permeability for 4-MUG is low. These data suggest that oral delivery of 4-MU is an effective way to achieve longer half-lives of the drug. Conversely, oral 4-MUG is not absorbed effectively.

In a study where mice were treated with 4-MU and 4-MUG for several weeks, we observed that mice that received 4-MU reached far higher drug levels than mice treated with 4-MUG. In this study we see a rapid increase in serum 4-MU as well as 4-MUG after 4 days of treatment. We also observed a plateau from that day on until the end of our measurements more than two weeks later. When we treated with 4-MUG and looked at the serum concentrations of 4-MU we saw a fast increase after one day which stayed at that relatively low level for the time of our experiment.

In a study where mice were treated multiple weeks with 4-MU and 4-MUG and the treatment was stopped, after cessation of therapy, we observed a rapid drop of 4-MU in the serum in the 4-MU treated mice. In those mice, the 4-MUG concentration reached a peak at one hour after treatment stop and was non-detectable at six hours. When mice were treated with 4-MUG and this was stopped, both 4-MU and 4-MUG were again detectable for less than six hours. These data indicate that drug persists in circulation for several hours after continuous dosing of 4-MU or 4-MUG.

While these studies illuminate the PK of 4-MU and 4-MUG in mice, the PK of 4-MUG in humans are not well studied. In healthy volunteers, the systemic exposure of 4-MUG after an i.v. dose was shown to be higher than that of 4-MU (28). Future clinical studies with 4-MU would benefit from a more thorough understanding of the PK of 4-MU itself and its metabolite 4-MUG.

We believe that our performed PK studies will aid the understanding and develop of dosing strategies including appropriate dose strength and frequency in animal models using 4-MU and 4-MUG. And hopefully these studies will ultimately better inform us about the use of 4-MU and maybe even 4-MUG in human clinical trials targeting HA synthesis in chronic and acute diseases (32).

## Abbreviations

4-MU: 4-methylumbelliferone
4-MUG: 4-methylumbelliferyl glucuronide

## Acknowledgements

This work was supported in part by the National Institutes of Health (NIH) grants R01 DK096087-01 and R01 HL113294-01A1 to P.L.B., U01 AI101984 to P.L.B, and grants from Stanford SPARK and the Stanford Children’s Health Research Institute (CHRI) to P.L.B. This work was also supported by grants from the Juvenile Diabetes Research Foundation (JDRF) 3-PDF-2014-224-A-N to N.N., and by a Pilot and Feasibility grant from the Stanford Diabetes Research Center, NIH P30DK116074 to N.N.

## Conflict of interest statement

N.N., G.K. and P.L.B. are listed as inventors of the patents-pending (PCT/US2014/050770, S17-131US/BLSU-1-65422, PCT/US2019/067911) filed by the Board of Trustees of the Leland Stanford Junior University.

## Contribution statement

N.N., M.A.U., S.V.M and P.L.B. conceptualization; N.N., G.K., P.L.B. supervision; N.N., and P.L.B. funding acquisition; N.N, G.K., N.L.H., A.H., M.A.U., J.R. and P.L.B investigation; N.N., G.K., N.L.H. and P.L.B. visualization; N.N., M.A.U. and G.K., methodology; N.N. and P.L.B. writing-original draft; N.N., G.K. and P.L.B. project administration.

## Data availability

The data are available from the corresponding author.

